# Broad PFAS binding with fatty acid binding protein 4 is enabled by variable binding modes

**DOI:** 10.1101/2025.01.10.632451

**Authors:** Aaron S. Birchfield, Faik N. Musayev, Abdul J. Castillo, George Zorn, Brian Fuglestad

## Abstract

Per- and polyfluoroalkyl substances (PFAS) are ubiquitous pollutants that bioaccumulate in wildlife and humans, yet the molecular basis of their protein interactions remains poorly understood. Here, we show that human adipocyte fatty acid-binding protein (FABP4) can bind a diverse array of PFAS, including next-generation replacements for legacy chemicals and longer-chain perfluorocarboxylic acids. Shorter-chain PFAS, although weaker binders, still displayed measurable affinities—surpassing those of their nonfluorinated analogs. We determined crystal structures of FABP4 bound to perfluorooctanoic acid (PFOA), perfluorodecanoic acid (PFDA), and perfluorohexadecanoic acid (PFHxDA), revealing three distinct binding modes. Notably, PFOA binds in two separate sites, and two distinct conformations define single-ligand binding of PFDA and PFHxDA. These arrangements enhance hydrophobic interactions within the binding cavity and likely explain the low micromolar dissociation constants observed in fluorescence competition assays. Our findings underscore the critical roles of chain length, headgroup functionality, and protein conformation in PFAS–FABP4 interactions. Given the emerging implications of the role of FABP4 in endocrine function, even subtle PFAS-induced perturbations could affect metabolic regulation and disease risk. Overall, this work highlights the value of direct structural and biochemical insights into PFAS–FABP4 interactions and paves the way for future research on PFAS transport and toxicological outcomes.

---

Per- and polyfluoroalkyl substances (PFAS) are a diverse class of synthetic chemicals characterized by fully or partially fluorinated carbon chains, which confer exceptional chemical and thermal stability and render them resistant to environmental degradation.^[1,2]^ These properties have facilitated the extensive use of PFAS in industrial and consumer products such as non-stick cookware, stain-resistant textiles, firefighting foams, and food packaging.^[3]^ Decades of widespread PFAS use have led to their global environmental distribution and measurable levels in wildlife and human populations.^[4,5]^ Understanding the mechanisms underlying PFAS bioaccumulation, transport, and interactions within biological systems is critical for assessing their potential health risks. Due to their chemical and physical similarity to lipids, PFAS have a propensity to interact with lipid carrier and lipid binding proteins, which are emerging as important targets to understand the health effects of PFAS exposure.^[6–8]^ To date, detailed structural investigation of PFAS protein binding in relation to has mostly focused on peroxisome proliferator-activated receptors (PPARs),^[9–11]^ human serum albumin (HSA),^[12]^ liver fatty acid-binding protein (FABP1),^[13–15]^ and transthyretin.^[16,17]^ All of these proteins are known lipid binders, leaving the role of other potential lipid binding protein interactions underexplored. Despite a great importance, at the time of writing, there exist only 11 experimentally determined PFAS bound structures available on the Protein Data Bank (PDB) (Table S1), which significantly limits understanding of the molecular drivers of PFAS-protein interactions.

The interactions of PFAS with FABP1 have been extensively studied.^[13,14,18]^ Interest in exploring this FABP isoform stems from the known accumulation of PFAS in the liver.^[19–22]^ However, it is possible that other FABP isoforms are likely effective binders of PFAS and responsible for bioaccumulation and health effects.^[23,24]^ Aside from liver and kidneys, PFAS are known to accumulate in circulation.^[25,26]^ While adipocyte fatty acid-binding protein (FABP4) is the only known secreted, circulatory fatty acid-binding protein, it has garnered surprisingly little attention in PFAS research, with an unrecognized role in human health effects from PFAS exposure. It is known as a fatty acid transporter in adipocytes,^[27–29]^ macrophages,^[30–32]^ and endothelial cells.^[27,33,34]^ Secreted, circulatory FABP4 was recently shown to regulate beta-cell function in pancreatic islets as a component of a hormone complex (Fabkin) that influences insulin secretion and metabolic homeostasis.^[35]^ While the role of ligand binding to FABP4 within the Fabkin complex has yet to be elucidated, potential for modulation by circulatory PFAS may explain observed correlations between PFAS exposure and diabetes, as well as related metabolic dysfunction.^[36–38]^ Elucidating the binding interactions between FABP4 and PFAS could provide new insights into biological transport and health effects of this interaction.

While PFAS such as perfluorooctanoic acid (PFOA) and perfluorooctanesulfonic acid (PFOS) have been phased out due to regulatory actions, accumulation in environmental reservoirs necessitates investigation into human health effects.^[39]^ Numerous replacement PFAS compounds, such as GenX, ADONA, and F-53B, are being introduced.^[40,41]^ Computational studies have characterized the protein binding and bioaccumulation potential of these emerging compounds, specifically in relation to HSA.^[42,43]^ However, experimental data on their interactions with other proteins, such as FABP4, as well as their environmental persistence and toxicity, remain limited. A wideranging understanding of protein binding to both legacy and emerging PFAS is thus essential to predict transport, accumulation patterns, and health effects across different classes of PFAS.

Here, we present the first comprehensive experimental study of PFAS interactions with FABP4. We measured the binding affinity for 16 PFAS by displacement of the fluorescent probe 8-anilinonaphthalene-1-sulfonic acid (ANS), establishing a relationship between chemical features of PFAS and their ability to bind with FABP4. We solved crystal structures of FABP4 bound to PFOA, perfluorodecanoic acid (PFDA), and perfluorohexadecanoic acid (PFHxDA), revealing distinct binding modes for each PFAS. These findings provide novel insights into the molecular mechanisms of PFAS binding to FABP4, which has high potential exposure to PFAS in its circulatory form. This study lays a foundation for understanding protein-driven PFAS bioaccumulation, transport, and toxicity in circulation.

Because FABP4 purified from *E. coli* is known to retain endogenous lipids^[44]^ that are challenging to remove,^[45]^ a butanol-extraction method was developed to achieve full delipidation at high yield (Details in the supplementary methods and Fig. S1). To understand the potential for PFAS interactions with FABP4, a comprehensive set of binding affinity measurements was obtained for a structurally diverse range of PFAS and alkanoic acid analogs of perfluorocarboxylic acids (PFCAs) (Figs. 1a, S2-5, Table S1). The results revealed clear evidence that the binding affinities of PFCAs are markedly dependent on chain length. Longer-chain PFCAs exhibit substantially higher binding affinities, as illustrated by the low micromolar K_d_ values of perfluorotetradecanoic acid (PFTeDA), perfluorotridecanoic acid (PFTrDA), and perfluorododecanoic acid (PFDoA), which all display K_d_ values below 1 μM. These data are consistent with extension of the hydrophobic perfluorocarbon tail enhancing the ability of PFAS to form stable hydrophobic interactions and more fully occupy the FABP4 lipid-binding cavity. Notably, this trend closely parallels the behavior of naturally occurring long-chain saturated fatty acids, such as palmitic acid, which display similar low micromolar K_d_ values.^[46,47]^

**Figure 1.**
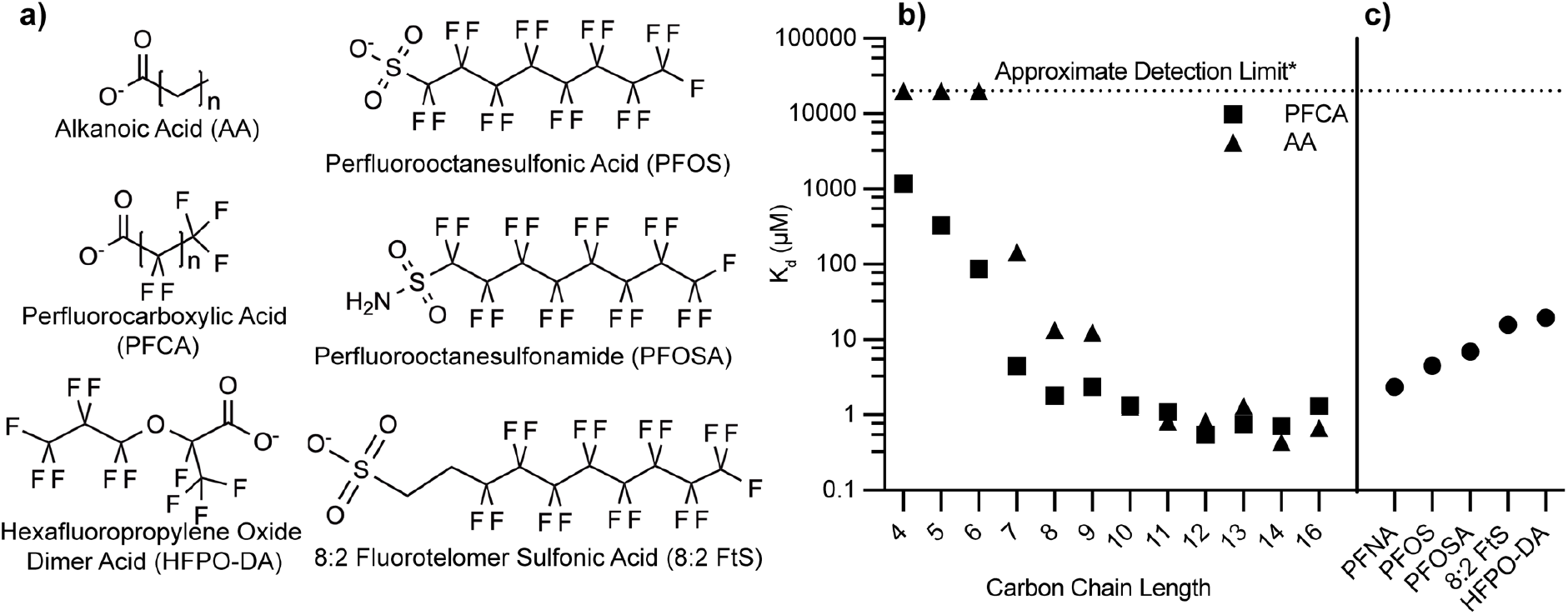
Structures and FABP4 binding affinities of compounds examined in this study. a) Chemical structures of PFAS and alkanoic acid compounds that were measured for affinity to FABP4. b) Binding affinity (K_d_) of perfluorocarboxylic acids compounds and their alkanoic acid analogs with FABP4. c) Affinity of PFAS compounds analogous to perfluorononanoic acid (PFNA) with alternate headgroups to FABP4. K_d_ values were converted from IC_50_ measurements obtained from ANS displacement assays. See supplement (Table S2) for detailed values including errors.

In contrast, shorter-chain PFCAs (e.g., perfluorohexanoic acid (PFHxA); perfluoropentanoic acid (PFPeA), perfluorobutanoic acid, PFBA) exhibited significantly weaker binding, with K_d_ values rising into the tens to hundreds of micromolar range. (Figs. 1B and S4, Table S1). This loss in binding strength can be attributed to diminished hydrophobic interactions and less extensive contacts formed within the FABP4 binding cavity. Nevertheless, even these shorter-chain PFAS exhibit measurable binding, which is especially pronounced in comparison to their non-fluorinated counterparts (Fig. S3). For example, perfluoroheptanoic acid (PFHpA) binds to FABP4 with a K_d_ of 4.46 ± 0.17 μM, representing an approximately 30-fold increase in affinity relative to its hydrogenated analog heptanoic acid (145 ± 3.1 µM). Similarly, while butanoic, pentanoic, and hexanoic acids displayed no detectable binding, their fluorinated analogs were readily observed to bind (Figs. 1, S3-4). These findings underscore a role for fluorination in stabilizing ligand-protein interactions, likely through enhanced hydrophobicity,^[48]^ even when hydrophobic chain length is insufficient to achieve strong binding in hydrogenated analogs.

In addition to chain length and the degree of fluorination, headgroup functionality exerts an influence on FABP4 binding affinity. This effect is evident when comparing perfluorononanoic acid (PFNA), PFOS, perfluorooctanesulfonamide (PFOSA), and 8:2 fluorotelomer sulfonic acid (8:2 FtS) (Fig. S4-5), each having eight fluorinated carbons, while 8:2 FtS has a longer chain, with ten carbons total, eight of which are fully fluorinated. Despite sharing a similarity in the number of fluorinated carbons, there are differences in binding affinity. PFNA binds more tightly to FABP4 than PFOS, PFOSA, or 8:2 FtS, which suggests that sulfonate and sulfonamide headgroups diminish affinity by interfering with stabilizing contacts typically achieved by carboxylate-containing ligands. Furthermore, the extended chain length of 8:2 FtS does not appear to confer any binding advantage, underscoring that headgroup identity and fluorination pattern are more critical determinants of affinity than additional methylene units. Together, these observations highlight the interplay between chain architecture, fluorination, and headgroup chemistry in governing PFAS-FABP4 interactions.

X-ray crystallographic analysis of FABP4 binding with PFOA, PFDA, and PFHxDA show distinct interactions that highlight the influence of chain length and fluorination on binding affinity (Fig. 2 and Table S1). Notably, PFOA was found to bind in two distinct sites within FABP4 (Fig. 2A). At the ‘primary site’, the carboxylate group is stabilized through hydrogen bonding with Arg 126 and Tyr 128, as well as a water-mediated hydrogen bond with Arg 106. The residues, Ala75, Thr29, Ala33, and Phe16 primarily interact through their hydrocarbon groups, contributing to hydrophobic interactions. PFOA binding at an observed ‘secondary site’ is stabilized by hydrogen bonding between its carboxylate group and Thr29 (Fig. 2A and S6). We note here that the assay likely reports only on the affinity of PFOA for binding to the primary site, due to the positioning of ANS here.^[49]^ Since the x-ray electron density is slightly less intense for the PFOA in the secondary site, we expect that binding is weaker here than for the primary site (Fig. S7).

**Figure 2.**
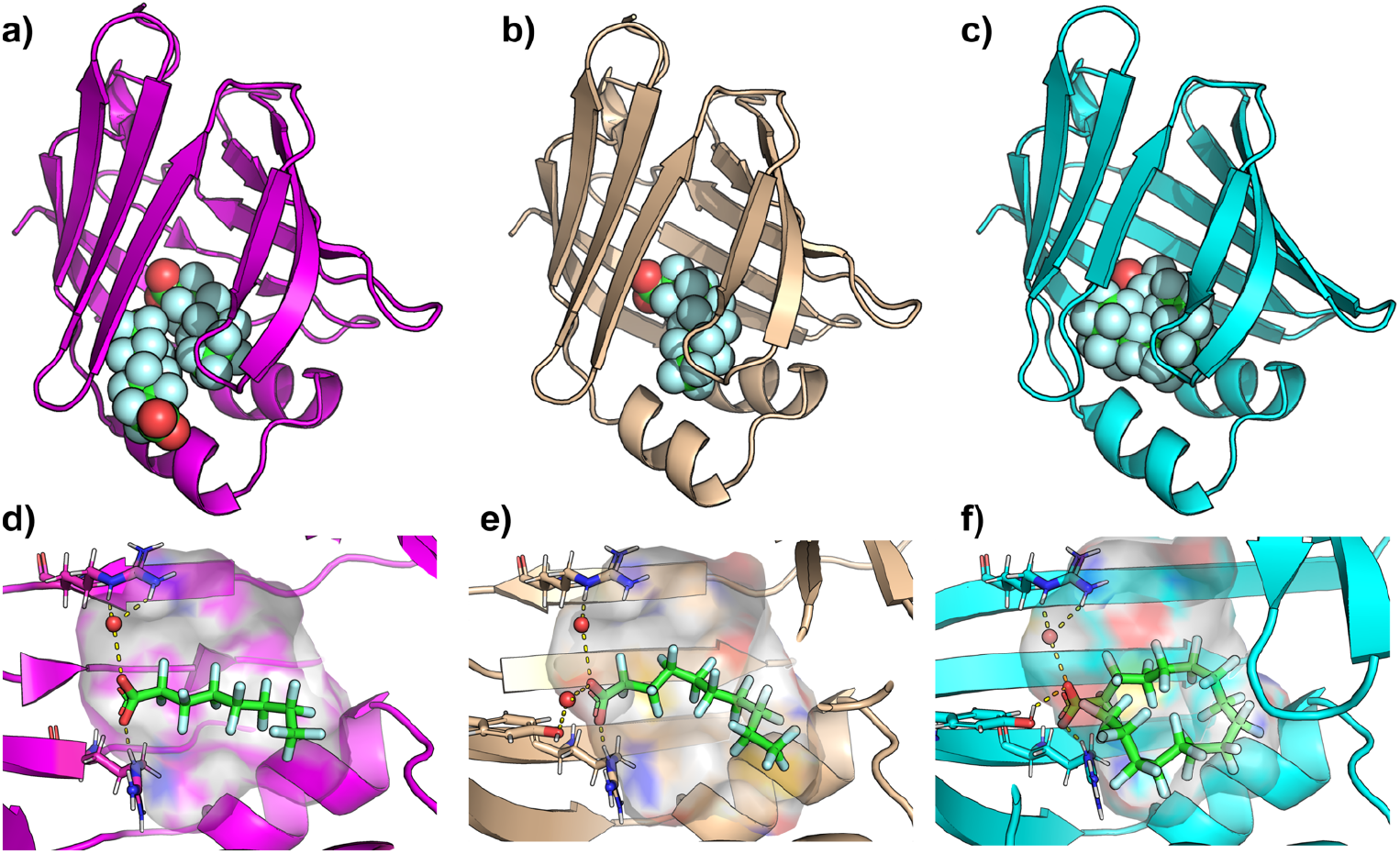
Crystal structures of PFOA, PFDA, and PFHxDA bound to FAPB4. a) X-ray crystallographic structure of PFOA (spheres) bound to FABP4 (magenta cartoon), displaying two PFOA molecules bound. b) X-ray crystallographic structure of FABP4 (wheat cartoon) with a single PFDA bound. c) X-ray crystallographic structure of FABP4 (cyan cartoon) with a single PFHxDA bound d) Zoomed image of the ‘primary’ PFOA, e) PFDA, or f) PFHxDA bound to the hydrophobic ligand binding cavity. The protein cavity is depicted as a semi-transparent surface, colored according to atomic identity: gray for hydrogens, blue for nitrogens, and red for oxygens, with carbons colored according to the cartoon coloring scheme. PFOA and PFNA are depicted in sticks, with key hydrogen-bonds and charge interactions highlighted in dashed yellow lines. For clarity, the water mediated H-bond between PFNA and S53 is omitted in panel d). Resolved water molecules that mediate PFAS-protein interactions are depicted in red spheres.

The binding pose of PFDA closely resembles that of the PFOA observed in the primary site (Fig. 2). In the PFDA structure, however, the extended fluorinated chain allows closer interactions with residues, Ala75, Asp76, Thr29, Ala33, and Phe16, reducing the distance between these residues and the ligand by approximately 1 Å compared to PFOA. (Figure 1 and S4). Both PFOA and PFDA induce an ‘open’ conformation of Phe57, which results in a larger hydrophobic cavity, compared to the closed conformation of Phe57 in the apo structure (Fig. 3 a-c). Interestingly, this residue is implicated in access of lipids to the hydrophobic cavity.^[44,50,51]^ The open conformation of Phe57 is necessary to accommodate the secondary PFOA ligand and the terminal fluoromethyl of PFDA. The extended tail length of PFDA requires a change in conformation compared to the apo state, which suggests that the ten-carbon chain induces energetically unfavorable steric clashes with the edge of the hydrophobic cavity. The small changes in PFDA proximity to cavity residues likely enhance hydrophobic interactions and van der Waals forces, compensating for the energetic cost of inducing a conformational change. We note that a similar conformational change is observed with PFOA but is induced by the presence of a second ligand. The PFHxDA crystal structure highlights distinct conformational features that accommodate its extended fluorinated chain. While PFDA remains fully extended, PFHxDA adopts a U-shaped conformation beginning around carbons 8–10, with carbon 9 forming the bend’s apex. This observation aligns with an *in-silico* analysis of rat FABP1, which suggested that PFCAs longer than ∼11 carbons must bend to fit within the binding pocket^[15]^ and resembles the binding mode of the hydrogenated analog, palmitic acid,^[44]^ a known natural ligand. Notably, Phe57 orients its aromatic ring inward in the PFHxDA-bound structure, resembling the closed conformation observed in the apo-FABP4, thereby enabling hydrophobic interactions (Fig. 3).

**Figure 3.**
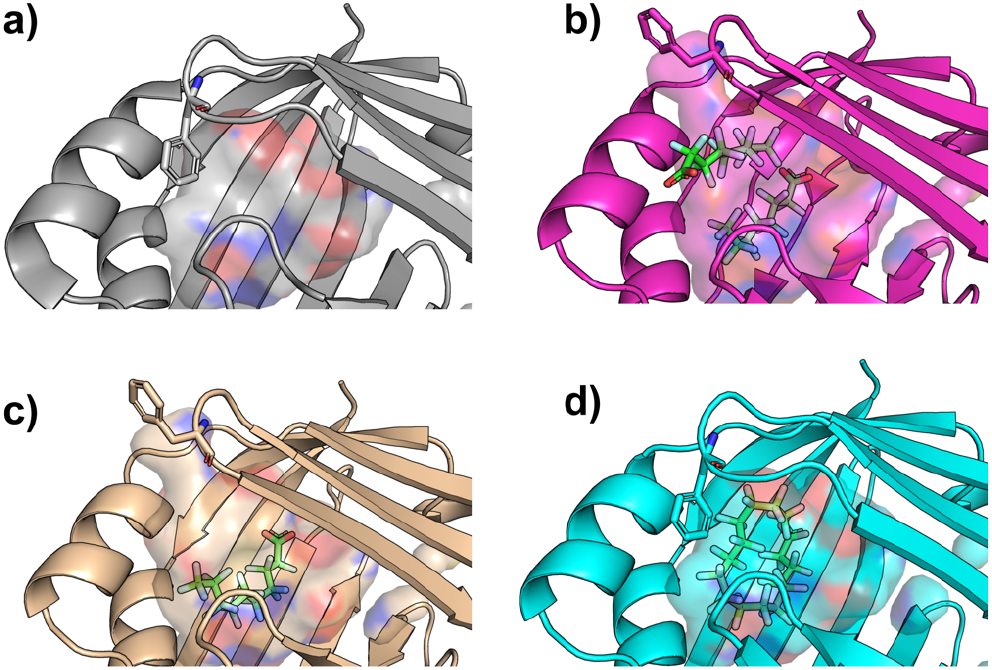
Phe57 changes conformation to accommodate PFAS ligands. Zoom of the region surrounding Phe57 in the crystal structures of a) apo-FABP4, b) PFOA-bound FABP4, c) PFDA-bound FABP4, and d) PFHxDA-bound FABP4. Phe57 and the PFAS ligands are displayed as sticks and the ligand binding cavity is displayed as a semi-transparent surface.

The crystal structures of PFOA, PFDA, and PFHxDA bound to FABP4 illustrate three distinct binding modes that underscore the protein’s adaptability to varying PFAS chain lengths. First, the PFOA-bound structure reveals two binding sites that accommodate shorter-chain PFAS. Second, the PFDA-bound structure showcases a fully extended perfluorocarbon tail, which induces an outward-facing Phe57, thereby creating sufficient space within the binding pocket. Third, the PFHxDA-bound structure adopts a U-shaped conformation, shifting Phe57 inward to maximize hydrophobic contacts. These observations, combined with our binding affinity data, indicate that while FABP4 can accommodate a range of PFCAs, those with longer chains must adopt a bent conformation to fit within the binding pocket. Inducing this degree of bending in a perfluorinated carbon chain requires more energy than for the corresponding hydrogenated carbon chain,^[15]^ which may be compensated for by the enhanced hydrophobicity from perfluorination compared to hydrogenated counterparts. Although a recent *in silico* docking study suggested that longer PFCAs cannot be accommodated within FABP4, ^[52]^ these findings highlight the importance of experimental starting points for computational predictions and suggest that the conformational flexibility of FABP4 plays a vital role in PFAS transport and bioaccumulation. Additionally, the structural consequences of PFAS binding reported here may support PFAS-binding protein design efforts.^[53]^

Our results indicate that FABP4 readily binds to a range of PFAS. This raises the possibility that PFAS exposure could, even at relatively low levels, interfere with normal function of FABP4 in lipid signaling and metabolism. The results underscore that the role of FABP4 in circulation and in the Fabkin hormone complex may be susceptible to modulation by PFAS, hinting at novel mechanistic links between PFAS exposure and metabolic disorders or diabetes. Future studies that evaluate the downstream consequences of PFAS–FABP4 binding – particularly under varying exposure scenarios – will be critical for understanding how these interactions might contribute to metabolic dysregulation or disease risk. These insights strengthen our grasp of PFAS-protein interaction mechanisms and lay important groundwork for further investigations into their systemic health impacts. ^[51]^

## Acknowledgements

We gratefully acknowledge Dr. Pingping Meng for helpful discussions. Research reported in this publication was supported by the National Institute of General Medical Sciences of the National Institutes of Health under Award Number R35GM147221 and Structural biology resources were provided by NIH Shared Instrumentation Grant S10OD021756. The content is solely the responsibility of the authors and does not necessarily represent the official views of the National Institutes of Health.

