## Supplemental Information for "Broad PFAS binding with fatty acid binding protein 4 is enabled by variable binding modes"

### Methods

#### Chemicals and Materials

Heptafluorobutyric Acid (Perfluorobutanoic Acid) was obtained from Sigma-Aldrich. Perfluoropentanoic Acid (PFPeA), Perfluorohexanoic Acid (PFHxA), Perfluorononanoic Acid (PFNA), Perfluorotetradecanoic Acid (PFTeDA), Perfluorotridecanoic Acid (PFTrDA), Perfluorododecanoic Acid (PFDoA), Perfluorodecanoic Acid (PFDA), Hexafluoropropylene Oxide Dimer Acid (HFPO-DA), Perfluorohexanesulfonamide (PFOSA), and Perfluorooctanesulfonic Acid (PFOS) were purchased from Cayman Chemical. Perfluoroheptanoic Acid and Perfluorooctanoic Acid (PFOA) were obtained from Sigma-Aldrich. Perfluorohexadecanoic Acid (PFHxDA) was acquired from Combi-Blocks.

Butanoic Acid, Hexanoic Acid, Heptanoic Acid, Octanoic Acid, Nonanoic Acid, Decanoic Acid, Dodecanoic Acid, Tetradecanoic Acid (myristic acid), and Hexadecanoic Acid (palmitic acid) were obtained from Sigma-Aldrich. Pentanoic Acid was sourced from Thermo Scientific. Undecanoic Acid and Tridecanoic Acid were acquired from Cayman Chemical. All other chemicals not explicitly mentioned in the text were purchased from commercial suppliers and used as received.

#### Protein Expression and Purification

Human Fatty Acid Binding Protein (FABP4) (Uniprot Accession: P15090) was recombinantly expressed in BL21(DE3) cells transformed with a pET-28a(+)-TEV vector and cultured in M9 media as described in Azatian et al.[1] Cells were pelleted by centrifugation at 5000g for 30 minutes at 4°C, then stored at -80°C or used immediately. Pelleted cells were resuspended in Lysis Buffer (50 mM Tris-Cl pH 7.5, 300 mM NaCl, 2 mM Dithiothreitol (DTT), 0.5 mM Ethylenediaminetetraacetic Acid (EDTA), 0.5% Triton X-100) containing cOmplete™ Mini EDTA-free Protease Inhibitor Cocktail (Millipore Sigma). Lysozyme (Gold Bio) was added to a final concentration of 0.2 mg/mL, and the suspension was incubated on ice for 30 minutes.

Cells were sonicated and centrifuged at 13,000g for 30 minutes at 4°C. The supernatant was collected, and imidazole was added to a final concentration of 10mM. The mixture was applied to a 2-mL nickel-nitrilotriacetic acid (Ni-NTA) resin column (Gold Bio) pre equilibrated with Equilibration Buffer (50 mM Tris-Cl pH 7.5, 300 mM NaCl, 10mM imidazole, and 2 mM DTT) using gravity flow. After collecting the flow-through, the resin was washed with 10 column volumes of Wash Buffer (50 mM Tris-Cl pH 7.5, 300 mM NaCl, 10-20 mM imidazole, and 2 mM DTT), or until protein concentration in the flow-through was negligible as determined by

Bradford assay. FABP4 was eluted with Elution Buffer (50 mM Tris-Cl pH 7.5, 300 mM NaCl, 300mM imidazole, and 2 mM DTT) and briefly desalted using a PD-10 desalting column equilibrated with 20 mM Tris-Cl pH 7.5, 100 mM NaCl, 2 mM DTT, and 0.5 mM EDTA.

Recombinant Tobacco Etch Virus (TEV) protease, expressed and purified in-house, was added to the desalted eluate at a ratio of 30 µg TEV protease per 1 mg FABP4 (determined by Bradford assay). Proteolytic digestion was performed overnight at 4°C. The digested sample was then applied to a fresh 2-mL Ni-NTA resin column, and the flow-through containing cleaved FABP4 was collected. The resin was washed with 5 mL of Equilibration Buffer and the wash fractions were combined with the flow-through. The purified, digested FABP4 sample was dialyzed overnight at 4°C against 1L of 20 mM Tris-Cl pH 7.5, 100 mM NaCl, 2 mM DTT, and 1mM EDTA. The dialyzed sample was concentrated to approximately 10mL in preparation for delipidation. Bradford assay measurements indicated typical yields of approximately 20 mg of FABP4, with SDS-PAGE analysis confirming the sample to be ≥95% homogeneous.

**FABP4 Delipidation**

FABP4 purified from *E. coli* is known to retain endogenous lipids,[2] which necessitates delipidation to prevent interference with downstream experiments. Previous studies have noted the difficulty of achieving complete delipidation of FABP4.[3] In our hands, methods such as lipidex-based extraction or guanidinium-induced denaturation and refolding resulted in ≥90% protein loss, yielding only 1–2 mg of purified protein. To address this, we developed a butanol-extraction method that achieved full delipidation while consistently yielding 10–12 mg of FABP4.

To delipidate the protein sample, n-butanol was used in a multi-step extraction process. Initially, 20% (v/v) n-butanol was added to the protein solution, which was then gently swirled and centrifuged at 4000 × g for 15 minutes at room temperature. The organic layer and interphase were carefully removed. This process was repeated twice using 15% (v/v) n-butanol, with centrifugation at 4000 × g for 10 minutes between each extraction. The organic layer and interphase were removed after each centrifugation. The interphase thickness was observed to decrease progressively with successive extractions. The solution was passed through a 0.22 µm filter to remove precipitates and lipid-protein complexes. Two additional extractions with 15% (v/v) n-butanol were performed, as described above, followed by a second filtration through a 0.22 µm filter. After this step, the interphase boundary between the water and n-butanol layers was observed to become less distinct. The delipidated protein solution was concentrated using an Amicon® Ultra 10kDa regenerated cellulose spin concentrator (Millipore). To prevent membrane swelling caused by n-butanol, the sample was concentrated by 50% increments, with an equal volume of Sample Buffer (20 mM Tris-Cl, pH 7.5, 100 mM NaCl, 2 mM DTT) added after each concentration step. Each step involved centrifugation at 3600 × g for 3–4 minutes. The protein solution was thoroughly mixed after each concentration step to prevent protein precipitation at the membrane. The protein was washed 6–8 times by diluting with Sample Buffer to 50% of the current volume and reconcentrating. This step effectively removed residual n-butanol from the protein solution. The concentrated protein solution was dialyzed against 1 L of Sample Buffer overnight at 4°C. Following dialysis, the protein was concentrated to 1mg/mL, aliquoted, and stored at -80°C. Protein quality and purity were assessed using SDS-PAGE with Coomassie staining and delipidation was confirmed using protein nuclear magnetic resonance (NMR) (Figure S1). Deuterated $H_2O$ ($D_2O$) was added to a 1-1.5 mg/mL sample of FABP4 at a final concentration of 10%, and $^{1}H$-$^{15}N$ HSQC spectra were collected to assess the lipidation state [3], using a Bruker AVANCE III 600 MHz instrument equipped with a room temperature QXI-probe. NMR data was processed with NMRPipe and analyzed with the NMRFAM-Sparky distribution.[4,5]

**Binding Inhibition Assay**

The binding affinity of 8-Anilinonaphthalene-1-sulfonic acid (ANS) (Cayman Chemical) with FABP4 was determined by mixing 0.5 uM ANS with increasing concentrations of FABP4 (0.01 uM-1 5uM) in 300 uL Assay Buffer (50 mM Tris-Cl pH 7.5, 100 mM NaCl, and 2 mM DTT). Samples were incubated for 2 minutes in the dark, and 250 µL of each sample was transferred to a Greiner UV-STAR® 96-well black microplate (Item No.: 655809). Fluorescence measurements were taken in triplicate using a SpectraMax ID5 microplate reader with an endpoint scan at an excitation wavelength of 370 nm and an emission wavelength of 480 nm. The data were analyzed in GraphPad Prism using nonlinear regression with a quadratic ligand depletion binding model, as described by the equation shown below:

$$Y = D_{max} * \frac{(K_d + x + p) - \sqrt{(K_d + x + p)^2 - 4px}}{2p} + BL$$

In this equation, $Y$ represents the observed fluorescence intensity, and $x$ corresponds to the ligand concentration, in this case, the FABP4 protein concentration (µM). The parameter $p$ is the fixed concentration of ANS held constant at 0.5 µM. $D_{max}$ represents the maximum fluorescence intensity observed when all binding sites are occupied, $K_d$is the apparent dissociation constant (µM), and $BL$ is the baseline fluorescence in the absence of ligand.

For the binding inhibition assay, a total of 275 µL of Assay Buffer containing 0.5 µM FABP4 and 0.5 µM ANS was mixed with 15 µL of a concentrated per- and polyfluoroalkyl substance (PFAS) or alkanoic acid stock solution. Each PFAS stock was prepared in ethanol or methanol, depending on the compound's solubility, and spanned a range of concentrations from 0.01 µM to 20,000 µM. Adding 15 µL of the stock to the assay tube resulted in the desired final ligand concentrations while maintaining a consistent final solvent concentration of 5% to increase solubility. For the 0 µM ligand control, 15 µL of the solvent (without ligand) was added to account for any minor reductions in fluorescence caused by the solvent itself. For the compound with the lowest reported solubility (PFHxDA, as indicated in the literature), the highest concentration tested was verified by dynamic light scattering (DLS) to confirm the sample was monodisperse and did not contain aggregates.[6] Samples were incubated for 2 minutes in the dark, and 250 µL of each sample was transferred to a Greiner UV-STAR® 96-well black microplate. Fluorescence measurements were taken in triplicate using a SpectraMax ID5 microplate reader with an endpoint scan at an excitation wavelength of 370 nm and an emission wavelength of 480 nm. The data were analyzed in GraphPad Prism using a sigmoidal four-parameter logistic (4PL) regression model, where $x$ represents the logarithm of the ligand concentration. The model was used to fit the experimental data and determine the $IC_{50}$ values for each tested compound, representing the concentration required to displace 50% of ANS bound to FABP4 (Figure S3-S5).

The $IC_{50}$ values obtained from the binding inhibition assay were converted to $K_i$ values using the $IC_{50}$-to-$K_i$ converter developed by Cer et al:[7]

$$K_i = \frac{I_{50}}{\frac{L_{50}}{K_d} + \frac{P_0}{K_d} + 1}$$

where $P_0$ and $L_{50}$ are free protein and ligand concentrations, respectively. The following parameters were entered into the converter: protein concentration of 0.5 µM, ligand (ANS) concentration of 0.5 µM, and the dissociation constant ($K_d$) of ANS for FABP4 as 0.508 µM, determined from the earlier assay (Figure S2). The $IC_{50}$ values for each inhibitor were used as the input parameter for inhibition. The calculations were performed using the 'free concentrations' option, which accounts for the unbound fractions of the protein and ligand in the system, and the $K_i$ values were taken from the output under the 'competitive' model. The resulting $K_i$ values represent the binding affinities of the tested PFAS compounds, accounting for the effects of protein and ligand concentrations on the measured $IC_{50}$ values. As the fluorescent probe ANS is well-characterized and binds in a 1:1 stoichiometry with FABP4,[8] the calculated $K_i$ values are reported as $K_d$ values to reflect the equilibrium dissociation constants for PFAS binding.

**Crystallization and Structure Determination of FABP4-Ligand complexes**

Complexes of FABP4 with PFOA, PFDA, and PFHxDA were prepared by addition of a 10-fold molar excess of the compound to a 1 mg/ml protein solution in 20mM Tris-HCl, pH 7.5, 100 mM NaCl, 2 mM DTT, and 0.01% Triton-100. Samples were incubated overnight at approximately 20°C with mild shaking. Samples were then concentrated to a final protein concentration of 8-10 mg/mL. Initial crystallization screenings were carried out with a Crystal Gryphon robot (Art Robbins Instruments). The following commercially available crystallization kits were used: SaltRx HT (Hampton Research) and Wizard Classic 1 and 2 blocks (Rigaku). Droplets were set up by mixing the protein-ligand complex sample (0.3 µl) with the crystallization reagent (0.3 µl) in a 1:1 ratio. They were then equilibrated against a 58 µl reservoir solution in 96-well sitting-drop INTELLI-PLATES. All plates were placed in a Gallery Plate Hotel at a constant temperature of 293K. Crystals appeared in one week under several conditions (K/Na phosphate, Na citrate tribasic, NH4 citrate and PEG-3000) at pH range 6.9-8.5. Attempts to improve the quality and the size of the crystals using the hanging-drop vapor-diffusion method were performed up to the microliter range in 24-well VDX crystallization plates (Hampton Research). Highly diffraction grade single crystals of FABP4-ligand complex obtained at pH 6.5 from 1.55-1.6 M Na citrate as a precipitant solution.

**Data collection, phasing, and model refinement**

For the x-ray data collection, crystals were flash-frozen in the liquid nitrogen stream mounted by nylon loops after dipping in a cryo-protectant solution for a few seconds (1.6 M Na citrate tribasic and 5% glycerol). Diffraction data was collected at 100 K up to 1.24 Å (FABP4-PFOA), 1.4 Å (FABP4-PFDA), and 1.3 5Å (FABP4-PFHxDA) resolutions on in-house copper rotating-anode generator (Rigaku MicroMax-007 HF) equipped with an EIGER R 4M detector. The X-ray data set was processed and scaled with *CrysAlis$^{Pro}$* 41.64.122a (Rigaku) and the CCP4 suite[9] of programs. Diffraction data for FABP4-PFOA and FABP4-PFDA complex crystals were indexed in space group C2 with one molecule in the asymmetric unit. For the FABP4-PFOA complex, unit cell dimensions were a = 118.752, b = 37.716, c = 28.546Å, β = 93.02° and for the FABP4-PFDA complex, a = 118.746, b = 37.709, c = 28.490Å, β = 92.50°. FABP4-PFHxDA complex crystals were indexed in space group P212121 with one molecule in the asymmetric unit, with unit cell dimensions a = 32.217, b = 53.564, and c =75.216Å. The structure was determined by molecular replacement using Phaser[10] from the PHENIX suite.[11] The starting model was the structure of the human adipocyte fatty acid-binding protein (PDB entry 7WC3). Auto model building resulted in an initial model of 129 amino-acid residues built with $R_{work}$ 18.89% and $R_{free}$ 20.54% (FABP4-PFOA complex), $R_{work}$ 23.54% and $R_{free}$ 29.23% (FABP4-PFDA complex), and $R_{work}$ 24.36% and $R_{free}$ 26.68% (FABP-PFHxDA complex). The structures were refined with the Phenix software along with manual model building using the graphics program Coot. The difference Fourier map revealed the presence of bound ligands in the active site of the FABP4. Ligand bound at the active site was clearly defined in the electron density map. Ligand was autofit into the electron density using the LigandFit procedure within Phenix, followed by manual rotation of CF2 groups using chi angles rotation mode in Coot.[12] The structures were refined to 1.24 Å resolutions with a final $R_{work}$ of 12.44% and $R_{free}$ of 15.70% (FABP4-PFOA complex), 1.4 Å resolution with a final $R_{work}$ of 13.96% and $R_{free}$ of 19.74% (FABP4-PFDA complex), and 1.35 Å resolution with a final $R_{work}$ of 17.81% and $R_{free}$ of 21.10% (FABP4-PFHxDa complex). Data collection and refinement statistics are summarized in Table S3.

**Table S1:** Experimentally Determined PFAS-Bound Protein Structures Available on the Protein Data Bank (PDB)

| Protein Name | PFAS Molecule | PDB Entry ID | Reference |
|---|---|---|---|
| Human Serum Albumin | Perfluorooctanesulfonic Acid (PFOS) | 4E99 | [13] |
| Human Serum Albumin | Perfluorooctanoic Acid (PFOA) | 7AAI | [14] |
| Human Serum Albumin | GenX | 7Z57 | [15] |
| Human Transthyretin | Perfluorooctanoic Acid (PFOA) | 5JID | [16] |
| Human Transthyretin | Perfluorooctanesulfonic Acid (PFOS) | 5JIM | [16] |
| Sea Bream Transthyretin | Perfluorooctanoic Acid (PFOA) | 6GON | [17] |
| Sea Bream Transthyretin | Perfluorooctanoic Acid (PFOA) | 6GOO | [17] |
| Human Heart Fatty Acid-Binding Protein | Perfluoroheptanoic Acid (PFHpA) | 7FD7 | (To be published) |
| Human Heart Fatty Acid-Binding Protein | Perfluorooctanoic Acid (PFOA) | 7FEK | (To be published) |
| Human Heart Fatty Acid-Binding Protein | Perfluorononanoic Acid (PFNA) | 7FEU | (To be published) |
| Human Peroxisome Proliferator-Activated Receptor Gamma (PPARγ) Ligand Binding Domain (LBD) | Perfluoraooctanoic Acid (PFOA) | 8U57 | [18] |

**Table S2:** Binding affinities of PFCAs, alkanoic acids, and PFAS with sulfonic acid, sulfonamide, ether, and alcohol headgroups. Data points represent triplicate measurements for each value; the $IC_{50}$ reported "±" values correspond to standard errors. The $K_i$ values were calculated as described in Cer et al.,[7] with their associated ± error propagated from $IC_{50}$ errors. The abbreviation "ND" indicates that no binding was detected.

| **C-chain** | **Perfluorocarboxylic Acids (PFCA)** | **$IC_{50}$ (μM) ± SD** | **$r^2$** | **$K_i$ (μM) ± error** |
|---|---|---|---|---|
| 4 | Perfluorobutanoic Acid (PFBA) | 2843 ± 140 | 0.9981 | 1181 ± 58 |
| 5 | Perfluoropentanoic Acid (PFPeA) | 792.8 ± 35.6 | 0.9972 | 329.2 ± 14.8 |
| 6 | Perfluorohexanoic Acid (PFHxA) | 208.9 ± 10.4 | 0.9996 | 86.65 ± 4.29 |
| 7 | Perfluoroheptanoic Acid (PFHpA) | 11.02 ± 0.44 | 0.9987 | 4.458 ± 0.178 |
| 8 | Perfluorooctanoic Acid (PFOA) | 4.669 ± 0.13 | 0.9994 | 1.82 ± 0.051 |
| 9 | Perfluorononanoic Acid (PFNA) | 5.944 ± 0.33 | 0.9980 | 2.35 ± 0.13 |
| 10 | Perfluorodecanoic Acid (PFDA) | 3.537 ± 0.09 | 0.9984 | 1.35 ± 0.034 |
| 11 | Perfluoroundecanoic Acid (PFUnDA) | 2.92 ± 0.12 | 0.9988 | 1.094 ± 0.045 |
| 12 | Perfluorododecanoic Acid (PFDoA) | 1.613 ± 0.057 | 0.9976 | 0.551 ± 0.020 |
| 13 | Perfluorotridecanoic Acid (PFTrDA) | 2.12 ± 0.072 | 0.9973 | 0.757 ± 0.026 |
| 14 | Perfluorotetradecanoic Acid (PFTeDA) | 2.002 ± 0.055 | 0.9981 | 0.7126 ± 0.020 |
| 16 | Perfluorohexadecanoic Acid (PFHxDA) | 3.446 ± 0.050 | 0.9987 | 1.312 ± 0.019 |
| **C-chain** | **Alkanoic Acids** | **$IC_{50}$ (μM) ± SD** | **$r^2$** | **$K_i$ (μM) ± error** |
| 4 | Butanoic Acid | ND | – | ND |
| 5 | Pentanoic Acid | ND | – | ND |
| 6 | Hexanoic Acid | ND | – | ND |
| 7 | Heptanoic Acid | 348.1 ± 7.47 | 0.9983 | 144.5 ± 3.10 |
| 8 | Octanoic Acid | 32.73 ± 1.72 | 0.9979 | 13.48 ± 0.71 |
| 9 | Nonoanoic Acid | 29.95 ± 0.61 | 0.9989 | 12.32 ± 0.25 |
| 10 | Decanoic Acid | 3.386 ± 0.136 | 0.9981 | 1.287 ± 0.052 |
| 11 | Undecanoic Acid | 2.249 ± 0.078 | 0.9978 | 0.8152 ± 0.028 |
| 12 | Dodecanoic Acid | 2.277 ± 0.072 | 0.9979 | 0.8268 ± 0.026 |
| 13 | Tridecanoic Acid | 3.459 ± 0.137 | 0.9985 | 1.318 ± 0.052 |
| 14 | Tetradecanoic Acid | 1.341 ± 0.055 | 0.9971 | 0.4381 ± 0.018 |
| 16 | Hexadecanoic Acid | 1.905 ± 0.082 | 0.9981 | 0.6723 ± 0.029 |
| **C-chain** | **PFAS with Sulfonic Acid, Sulfonamide, Ether, and Alcohol Headgroups** | **$IC_{50}$ (μM) ± SD** | **$r^2$** | **$K_i$ (μM) ± error** |

| | | | | |
|---|---|---|---|---|
| 8 | Perfluorooctanesulfonic Acid (PFOS) | 11.13 ± 0.53 | 0.999 | 4.504 ± 0.214 |
| 8 | Perfluorooctanesulfonamide (PFOSA) | 17.02 ± 0.93 | 0.998 | 6.95 ± 0.38 |
| 6 | Hexafluoropropylene Oxide Dimer Acid (HFPO-DA) | 47.32 ± 2.65 | 0.998 | 19.53 ± 1.09 |
| 10 | 8:2 Fluorotelomer Sulfonic Acid (8:2 FtS) | 38.35 ± 2.04 | 0.999 | 15.81 ± 0.84 |
| 10 | 1H,1H,2H,2H-Perfluoro-1-decanol (8:2 FTOH) | ND | – | ND |

**Table S3:** X-Ray crystal data collection and refinement statistics.

| | FABP4-PFOA | FABP4-PFDA | FABP4-PFHxDA |
|---|---|---|---|
| Space group | C2 | C2 | P 212121 |
| Unit-cell *a, b, c* (Å) | 118.752, 37.716, 28.546, β = 93.02(°) | 118.746, 37.709, 28.490 β = 92.50(°) | 32.217, 53.564, 75.216 |
| Resolution (Å)<br>Total reflections | 22.51–1.24 (1. 26 – 1.24)<br>505153 (2913) | 27.29–1.40 (1.42 – 1.40)<br>222985 (6098) | 27.61-1.35 (1.37-1.35)<br>250919 (7717) |
| Unique reflections | 35429 (1325) | 24826 (1205) | 29408 (1430) |
| Redundancy | 14.3 (2.2) | 9.0 (5.1) | 8.5 (5.4) |
| Completeness (%) | 98.7 (77.3) | 99.4 (95.9) | 100.0 (100.0) |
| Average I/σ(I) | 52.8 (6.9) | 33.7 (1.9) | 25.5 (2.2) |
| $R_{merge}$ (%)[a] | 2.5 (8.3) | 8.0 (77.3) | 4.6 (79.7) |
| Refinement Statistics | | | |
| Resolution (Å) | 22.51-1.24 (1.27-1.24) | 22.46-1.40 (1.46-1.40) | 27.61-1.35 (1.39-1.35) |
| No. of reflections | 35428 (2266) | 24819 (2643) | 29348 (2623) |
| $R_{work}$ (%) | 12.44 (12.44) | 13.96 (20.49) | 17.81 (20.88) |
| $R_{free}$ (%)[b] | 15.70 (16.89) | 19.74 (29.22) | 21.10 (23.81) |
| R.m.s.d. bonds (Å) | 0.006 | 0.006 | 0.005 |
| R.m.s.d. angles (°) | 1.114 | 0.950 | 1.099 |
| Dihedral angles | | | |
| Most favored (%) | 98.5 | 99.2 | 96.9 |
| Allowed (%) | 1.5 | 0.8 | 3.1 |
| Average B ($Å^2$) / atoms | | | |
| All atoms | 13.62 | 18.70 | 21.57 |
| Protein<br>Ligand<br>Solvent<br>Number of non-hydrogen atoms | 10.12<br>34.13<br>25.49<br>1399 | 15.41<br>67.64<br>28.36<br>1334 | 13.70<br>157.24<br>28.51<br>1289 |
| Macromolecules<br>Ligands<br>Water | 1108<br>50<br>241 | 1089<br>31<br>214 | 1030<br>49<br>210 |
| Protein residues | 134 | 134 | 131 |

[a]$R_{merge} = \Sigma_{hkl}\Sigma_i|I_i(hkl) - \langle I(hkl)\rangle|/\Sigma_{hkl}\Sigma_i I_i(hkl)$. [b]$R_{free}$ was calculated from 5% randomly selected reflection for cross-validation. All other measured reflections were used during refinement.

### Supplemental Figures

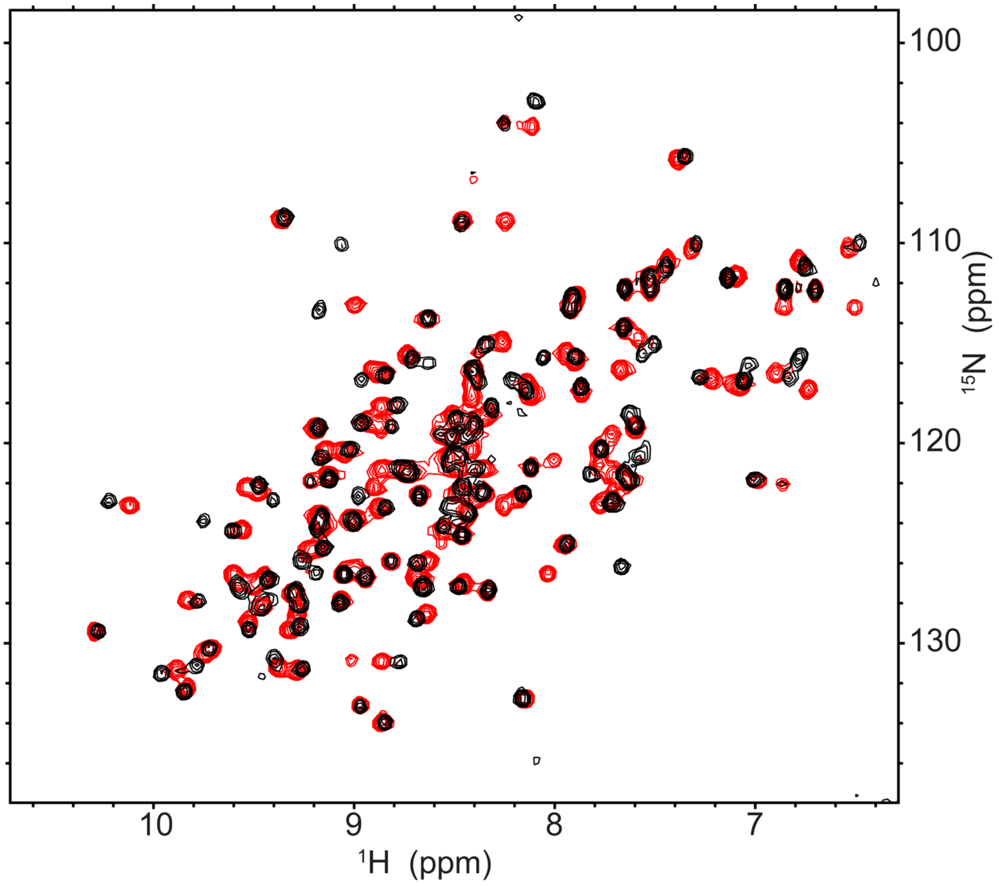


**Figure S1.** $^{1}$H-$^{15}$N HSCQ Spectra of human FABP4 before (black) and after (red) delipidation by butanol extraction. Spectra were collected at 25° with a 600 MHz instrument.

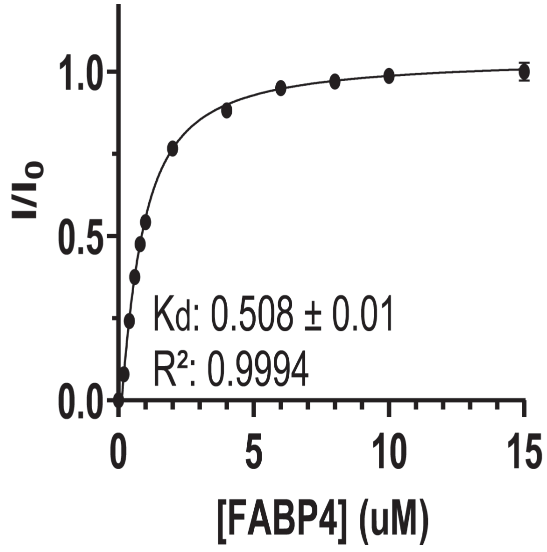


**Figure S2.** Binding Affinity of ANS for FABP4. Graph showing the normalized fluorescence intensity of FABP4 (0.01 μM to 15 μM). Data points represent triplicate measurements, with error bars indicating the standard deviation. Nonlinear regression was performed using a Quadratic Ligand Depletion Binding Model, yielding a dissociation constant ($K_d$) of 0.508 μM for the FABP4-ANS interaction.

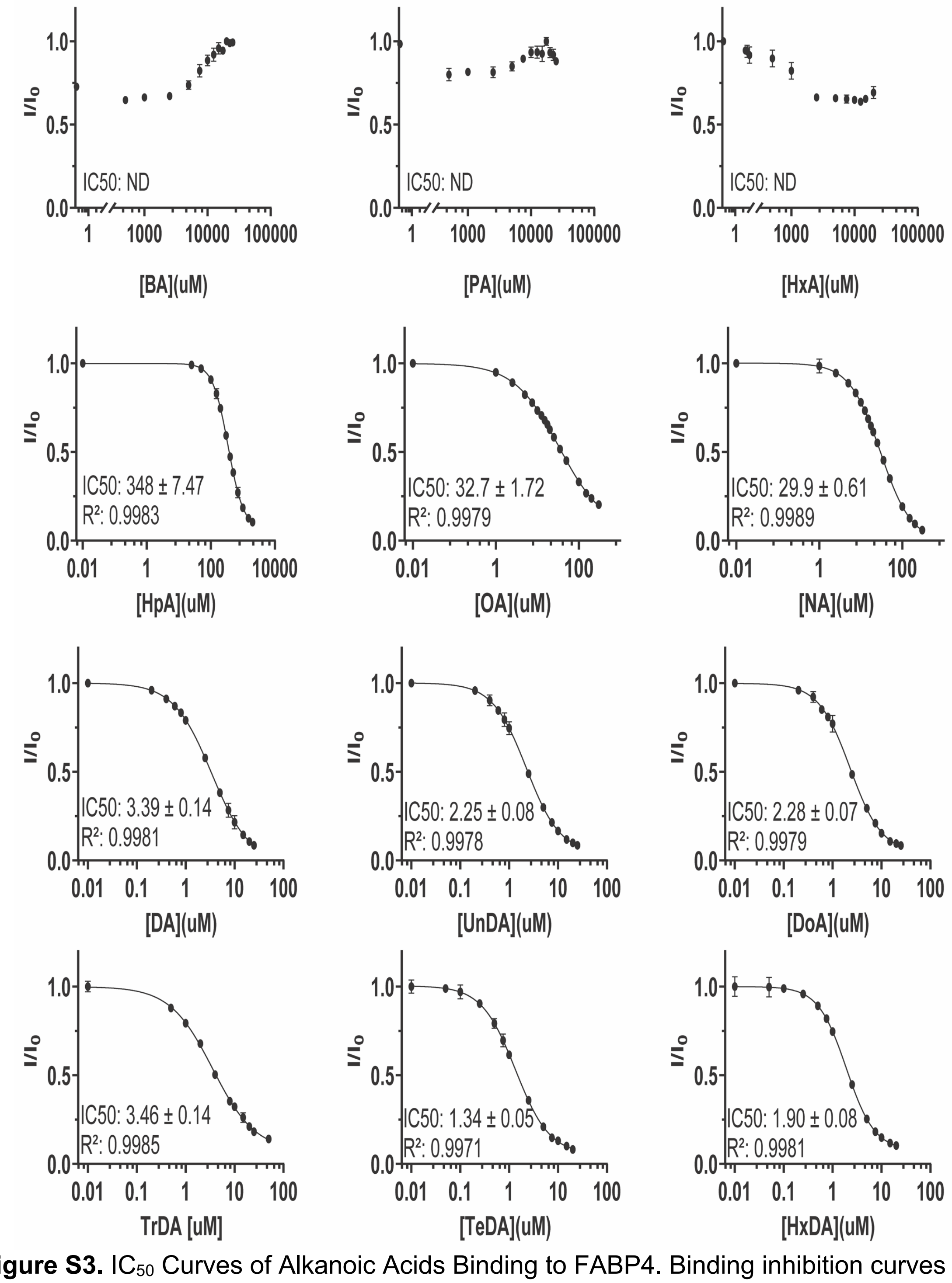


**Figure S3.** $IC_{50}$ Curves of Alkanoic Acids Binding to FABP4. Binding inhibition curves of alkanoic acids tested for displacement of ANS from FABP4. Each curve represents the normalized fluorescence intensity relative to the control (0 μM ligand), plotted against the log concentration of the ligand. $IC_{50}$ values were determined using a four-parameter logistic (4PL) regression model in GraphPad Prism. Data points represent the mean of triplicate measurements, and error bars indicate the standard deviation.

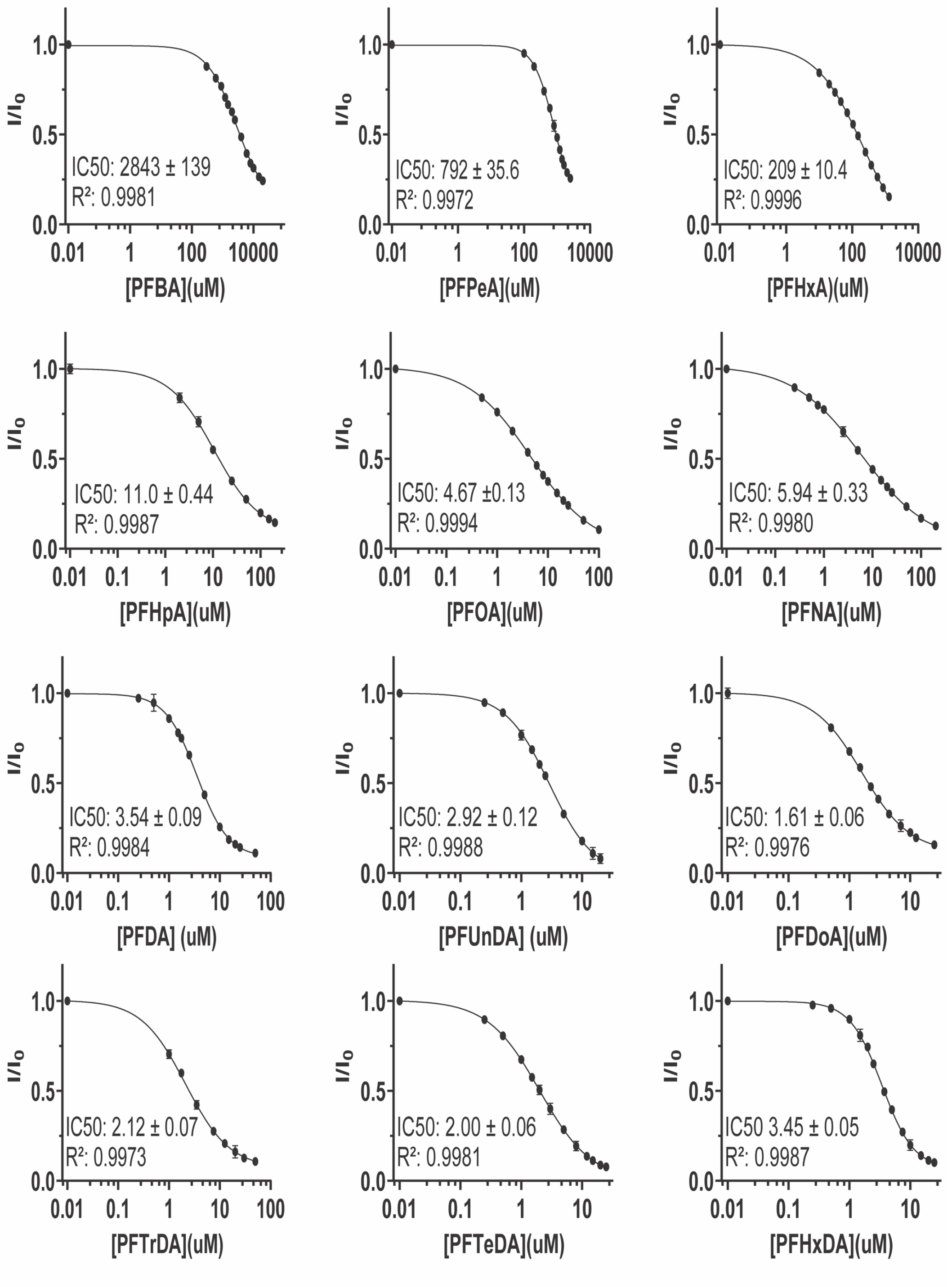


**Figure S4.** $IC_{50}$ Curves of Perfluorinated Carboxylic Acids (PFCAs) Binding to FABP4. Binding inhibition curves of PFCAs tested for displacement of ANS from FABP4. The curves display the normalized fluorescence intensity relative to the control (0 μM ligand) plotted against the log concentration of the ligand. $IC_{50}$ values were calculated using a 4PL regression model. Data represent triplicate measurements, with error bars showing the standard deviation.

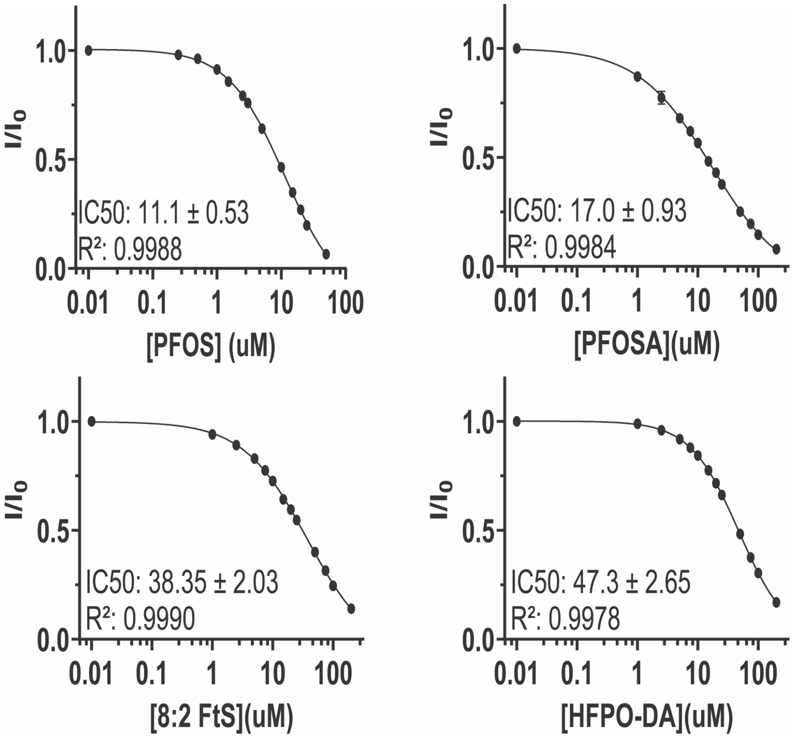


**Figure S5.** $IC_{50}$ Curves of Other PFAS Compounds Binding to FABP4. $IC_{50}$ curves for the displacement of ANS from FABP4 by additional PFAS compounds, including perfluorooctanesulfonic acid (PFOS), perfluorooctanesulfonamide (PFOSA), hexafluoropropylene oxide dimer acid (HFPO-DA), and 8:2 fluorotelomer sulfonic acid (8:2 FtS). Normalized fluorescence intensity relative to the control (0 μM ligand) is plotted against the log concentration of each compound. $IC_{50}$ values were determined using a 4PL regression model, with data shown as the mean of triplicate measurements and error bars indicating standard deviation.

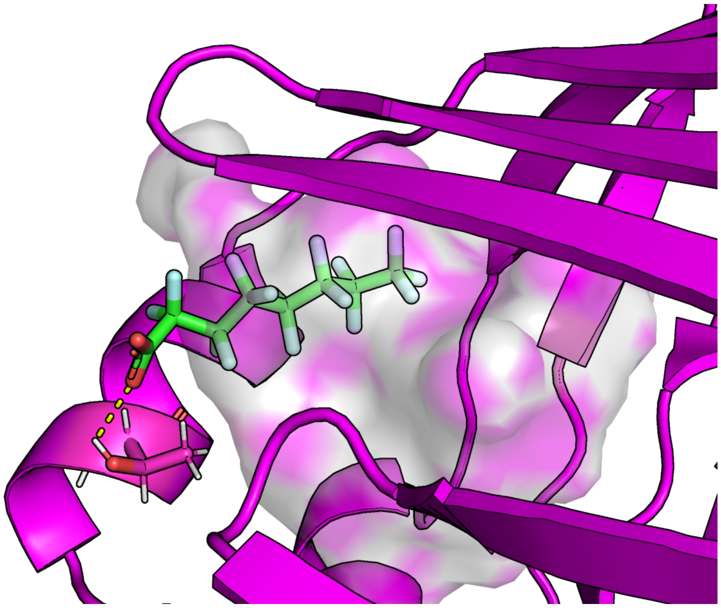

**Figure S6.** Zoom of the crystal structures of PFOA bound to FAPB4 in the secondary binding site. The protein cavity is depicted as a semi-transparent surface, colored according to atomic identity: gray for hydrogens, blue for nitrogens, and red for oxygens, with carbons magenta. PFOA and Thr29 are depicted as sticks, with their hydrogen-bond highlighted with a dashed yellow line. For clarity, PFOA bound in the primary site is omitted.

a)

b)

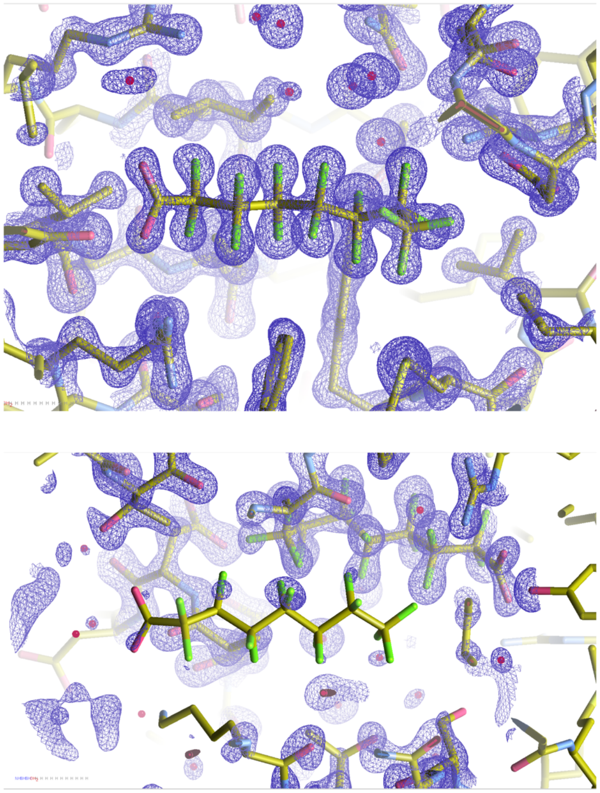

**Figure S7.** Electron density 2Fo − Fc maps with contour level σ = 2.0 of PFOA in the (a) primary and (b) secondary binding sites.

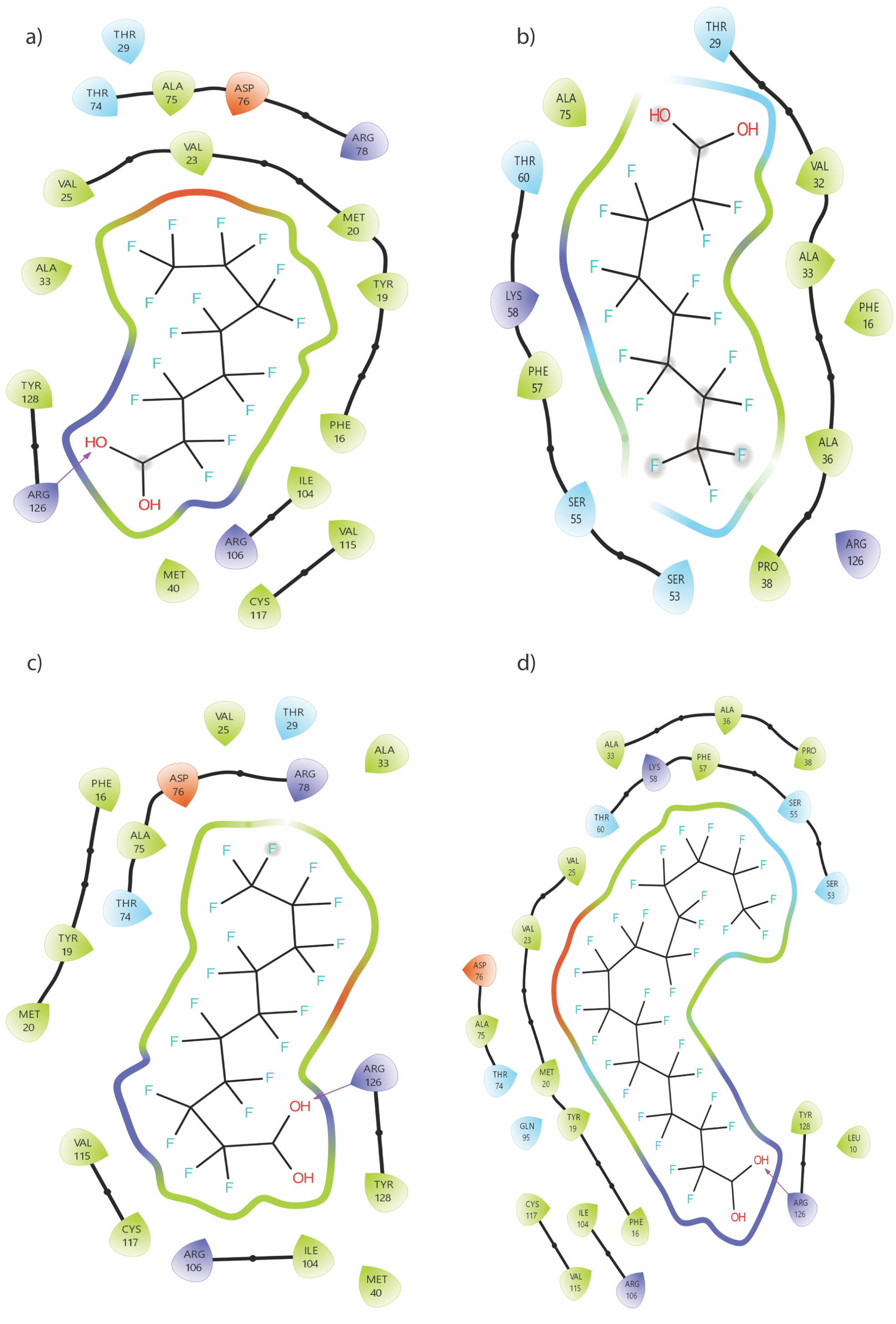


**Figure S8:** Ligand interaction diagrams of FABP4 with PFOA (Site 1) (a), PFOA (Site 2) (b), PFDA (c), and PFHxDA (d). Residues within 4 Å of the ligand include hydrophobic residues (green), positively charged residues (purple), negatively charged residues (red), and polar residues (light blue). Hydrogen bonds are indicated by purple arrows, and black lines denote residues on the same structural element (loop, β-sheet, or α-helix). Images were generated using Maestro Viewer.[19]
